## Supplementary Infomation for "Optogenetic activation of dorsal raphe serotonin neurons induces brain-wide activation"

### Supplementary Methods

#### Dataset

An optogenetic functional MRI dataset was acquired from openneuro (Project_ID: Mouse_opto_DRN, https://doi.org/10.18112/openneuro.ds001541.v1.1.2, for details) [1, 3]. The dataset comprises functional MRI experimental data with ePET-Cre mice expressing ChR2-YFP or YFP in dorsal raphe nucleus 5-HT neurons. Images were taken with cerebral blood volume or BOLD fMRI with a 7 T Pharmascan scanner of mice anesthetized with 0.5% isoflurane + 0.2mg/kg/h s.c. medetomidine. Anatomical images were made with a gradient echo FLASH sequence with the following parameters: 120 x 120 matrix, 20x17.5 $mm^2$ field of view, repetition time (TR)/effective echo time (TE) 1500/1.97 (ms), 14 coronal slices, slice thickness: 0.5 mm, slice gap 0.15 mm.

After anatomical images were taken, CBV images were further executed with multi-shot gradient echo EPI with the following parameters: 2 segments, 64 x 64 matrix, 20x17.5 $mm^2$ field of view, TR/TE 1000/5.6 (ms), flip angle: 90° , bandwidth:250k(Hz), 14 coronal slices, slice thickness: 0.5 mm, slice gap 0.15 mm, and 360 or 720 repetitions for the short-block and long-block protocols, respectively.

Laser stimulation protocols were composed of ON/OFF stimulation patterns. 20 Hz stimulus trains for 20 seconds were delivered 6 times per min in the short-block and every 3 min in the long block.

63 runs with the short-block protocol with laser power (473 nm, 4-40 mW) from eight mice were used for the subsequent analysis.

**Preprocessing and de-noising for the CBV fMRI dataset**

Similar preprocessing was applied to images (see "Preprocessing and de-noising" in Methods). 10-fold magnification of images was accomplished with SPM12. Then, the voxel size was 2 x 2 x 3 mm in analytical steps. Pre-processing, including motion correction, re-alignment, co-registration, normalization to the C57BL6/J template, and spatial smoothing (kernel with 4 x 4 x 6 mm) were subsequently applied with SPM12.

After pre-processing, the general linear model in SPM12 was executed to map activation patterns.

**General Linear Model (GLM) and beta values with CBV fMRI dataset**

A general linear model (GLM) for this study was executed with preprocessed CBV images to map beta values of parameter estimation. For statistical functional mapping of CBV images, each voxel under uncorrected P value < 0.001 was thresholded, and with FWE by thresholding pFWE < 0.05 with pTFCE. Statistical second-level, voxel-wise analysis was subsequently performed.

Regional beta values were extracted from 28 in-house ROIs based on the Allen Brain Atlas (see "Region of Interest" in Methods).

**Extraction of regional BOLD signal changes and peak intensities**

Each regional CBV signal change (ΔCBV) was calculated by averaging regional time series on voxels, after first aligning them with each stimulation timing from 28 ROIs based on the Allen Brain Atlas (see "Regions of Interest" in Methods) [3]. Average ΔCBV time series were detrended with signal means. Regional time series were further divided by onset of each stimulation timing, and each reginal peak intensity was further calculated with the max of ΔCBV signals in each 60-sec stimulation cycle.

| Software List | | | | |
|---|---|---|---|---|
| IDs | Name | Contributor | Purpose | URLs |
| 1 | Statistical Parametric Mapping 12(SPM12) | The Wellcome Centre for Human Neuroimaging, UCL | MRI data analysis | https://www.fil.ion.ucl.ac.uk/spm/ |
| 2 | MarsBaR region of interest toolbox for SPM | | Brain Reference Registration | http://marsbar.sourceforge.net/ |
| 3 | MRIcroGL | Chris Rorden, Neuropsychology Lab, University of South Carolina | 3D visualization of MRI data | https://www.nitrc.org/ projects/mricrogl/ |
| 4 | Probabilistic Threshold-free Cluster Enhancement (pTFCE) [5] | Tamás Spisák | A cluster-enhancement method | https://spisakt.github.io/pTFCE/ |
| 5 | Allen Brain Atlas | Allen Brain Institutes | Reference atlas, 5HT projection map, 5HT receptor gene expressions | https://portal.brain-map.org/ |
| 6 | Templates for In Vivo Mouse Brain [2] | Keigo Hikishima (k-hikishima@ aist.go.jp) | Mouse brain templates for registration and normalization | https:// www.nitrc.org/ projects/tpm_mouse |
| 7 | Gramm [4] | Pierre Morel | Visualization of Violin Plotting | https://github.com/piermorel/gramm |
| 8 | in-house analysis programs | Neural Computational Unit | Analysis for time series, Behavioral analysis | https://github.com/Taiyou/OptofMRI_5HT<br>https://doi.org/10.5281/zenodo.10956557 |
| 9 | MATLAB | MathWorks | Main analyses and data visualization | https://mathworks.com/ |
| 10 | A CBV opfMRI dataset | Joanes Grandjean | Analysis for functional maps | https://doi.org/10.18112/openneuro.ds001541.v1.1.2<br>https://doi.org/10.34973/raa0-5z29 |

Supplementary Table S1. Software employed in this study

| ROI id | ROI name | Allen Brain Atlas Name |
|---|---|---|
| 1 | Frontal polar area (FP) | Forntaol pole, layer 1, 2/3, 5, 6a, 6b |
| 2 | Orbital area (OFC) | Orbital area, lateral/medial/ventral/ventrolateral part, layer 1, 2/3, 5, 6a, 6b |
| 3 | Medial prefrontal cortex (mPFC) | Prelimbic area, layer 1, 2, 2/3, 5, 6a, 6b & Infralimbic area, layer 1, 2, 2/3, 5, 6a, 6b |
| 4 | Secondary motor area (M2) | Secondary Motor area, layer 1, 2/3, 5, 6a, 6b |
| 5 | Anterior cingulate cortex (ACC) | Anterior cingulate area, layer 1, 2/3, 5, 6a, 6b & Anterior cingulate area, dorsal/ventral part, layer 1, 2/3, 5, 6a, 6b |
| 6 | Primary motor area (M1) | Primary motor area, layer 1, 2/3, 5, 6a, 6b |
| 7 | Supplementary somatosensory area (SS) | Supplemental somatosensory area, layer 1, 2/3, 4, 5, 6a, 6b |
| 8 | Ventral retrosplenial cortex (vRSC) | Retrosplenial area, ventral part, layer 1, 2/3, 4, 5, 6a, 6b |
| 9 | Dorsal retrosplenial cortex (dRSC) | Retrosplenial area, dorsal part, layer 1, 2/3, 4, 5, 6a, 6b |
| 10 | Temporal association area (TAA) | Temporal association areas, layer 1, 2/3, 4, 5, 6a, 6b |
| 11 | Primary auditory area (A1) | Primary auditory are, layer 1, 2/3, 4, 5, 6a, 6b |
| 12 | Primary somatosensory area (S1) | Somatosensory area, layer 1, 2/3, 4, 5, 6a, 6b, Primary somatosensory area, layer 1, 2/3, 4, 5, 6a, 6b, Primary somatosensory area, nose/barrel field, layer 1, 2/3, 4, 5, 6a, 6b |
| 13 | Posterior parietal association area (PPA) | Posteior parietal association areas, layer 1, 2/3, 4, 5, 6a, 6b |
| 14 | Primary visual area (V1) | Primary visual area, layer 1, 2/3, 4, 5, 6a, 6b |
| 15 | Nucleus accumbens (NAc) | Nucleus accumbens |
| 16 | Caudate putamen (CPu) | Caudoputamen |
| 17 | Bed nucleus of steria terminalis (BST) | Bed nuclei of the stria terminalis, anterior/posterior division |
| 18 | Globus pallidus external part (GPe) | Globus pallidus, external segment |
| 19 | Globus pallidus internal part (GPi) | Globus pallidus, internal segment |
| 20 | Thalamic reticular nucleus (RT) | Reticular nucleus of the thalamus |
| 21 | Ventral pallidus (VP) | Pallidum, ventral region |
| 22 | Dentate gyrus of hippocampus (DG) | Dentate gyrus, molecular/polymorph/granule cell layer/subgranular zone, Dentate gyrus crest, molecular/polymorph/granule cell layer, Dentate gyrus lateral/medial blade, molecular/polymorph/granucle cell layer |
| 23 | CA1 of hippocampus (CA1) | Field CA1, stratum lacunosum-moleculare/stratum oriens/pyramidal layer/stratum radiatum |
| 24 | Lateral habenula (LHb) | Lateral habenula |
| 25 | Substantia nigra pars reticulata (SNr) | Substantia nigra, reticular part |
| 26 | Ventral tegmental area (VTA) | Ventral tegmental area |
| 27 | Medial raphe nucleus (MRN) | Superior central nucleus raphe, lateral/medial part |
| 28 | Dorsal raphe nucleus (DRN) | Dorsal nucleus raphe |

Supplementary Table ST2. Correspondence between ROIs and names in the Allen Brain Atlas

| | Session 1 | Session 2 | Session 3 | Session 4 |
|---|---|---|---|---|
| $R^2$ statistics | 0.5851 ± 0.1574 | 0.5505 ± 0.0863 | 0.5240 ± 0.1166 | 0.5437 ± 0.1100 |
| F statistics | 5.1849 ± 3.9732 | 3.6719 ± 1.2474 | 3.4689 ± 1.6842 | 3.7058 ± 1.6093 |
| P value | 0.0506 ± 0.0774 | 0.0352 ± 0.0464 | 0.0599 ± 0.0609 | 0.0468 ± 0.0570 |
| EEV | 0.1252 ± 0.0644 | 0.1009 ± 0.0224 | 0.1316 ± 0.0994 | 0.0630 ± 0.0259 |

Supplementary Table ST 3. Multiple linear regression with beta values from Sessions1-4. Multiple linear regression was applied to beta values from each subject for all sessions. Each value indicates the mean ± standard error of mean.

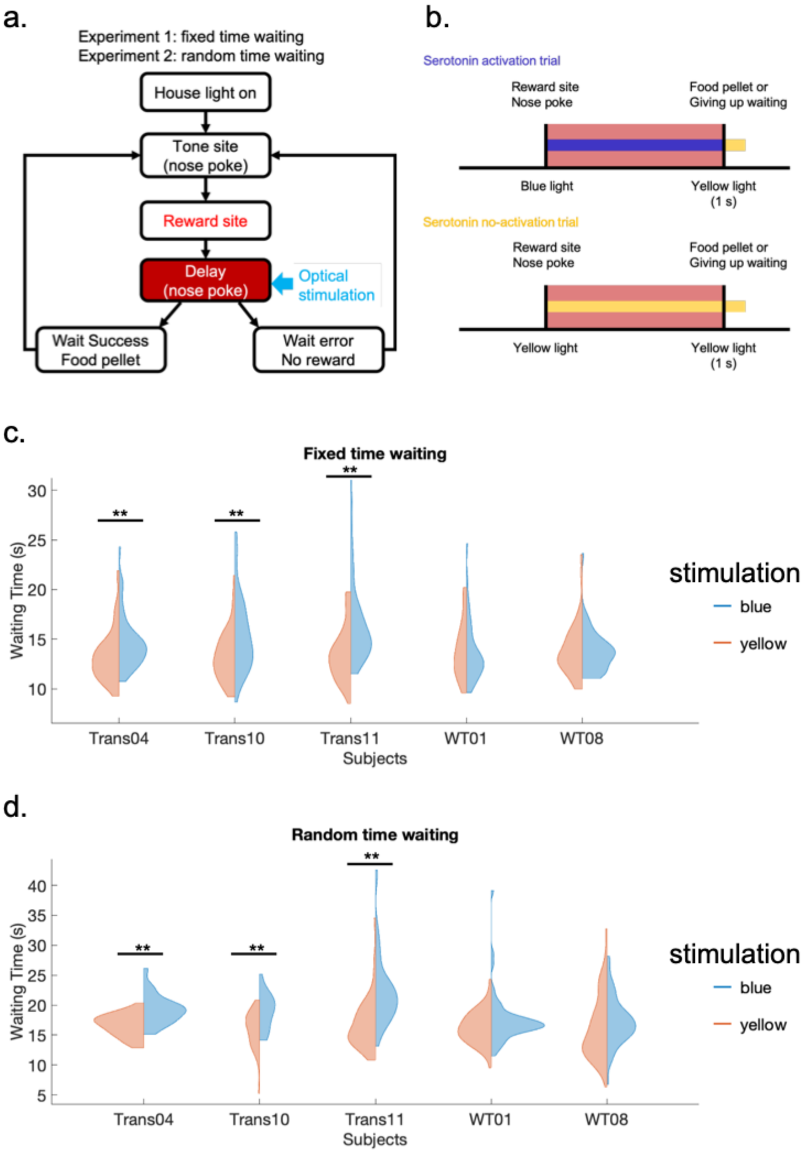


Supplementary Figure S1. Waiting Task. (a) Sequence of the tone-food waiting task. (b) Time sequence of optogenetic illumination. In serotonin activation trials, blue light stimulation was delivered until a trial ended with delivery of a food pellet or quitting. In serotonin no—activation trials, yellow light stimulation was delivered until the trial ended. After the trial ended, 1 s of yellow light was delivered. Blue and orange color indicate blue and yellow light stimulation. Red regions indicate reward delay periods. (c) Experiment 1: Fixed-time waiting (6.0 s). In this experiment, the reward delay duration for a success trial was fixed at 6 s. For an omission trial, no food reward was given. ** $p_{\text{Bonferroni-corrected}} < 0.01$ using the Wilcoxon rank sum test with Bonferroni correction. Blue and orange regions indicate distributions of blue light and yellow light stimulation. (d) Experiment 2: Random time waiting (2.0, 6.0, or 10.0 s). In this experiment, the reward delay duration for a successful trial was randomly delivered after 2, 6, or 10 s. For an omission trial, mice received no food reward. ** $p_{\text{Bonferroni-corrected}} < 0.01$ using the Wilcoxon rank sum test with Bonferroni correction. Blue and orange regions indicate distributions of blue light and yellow light stimulation.

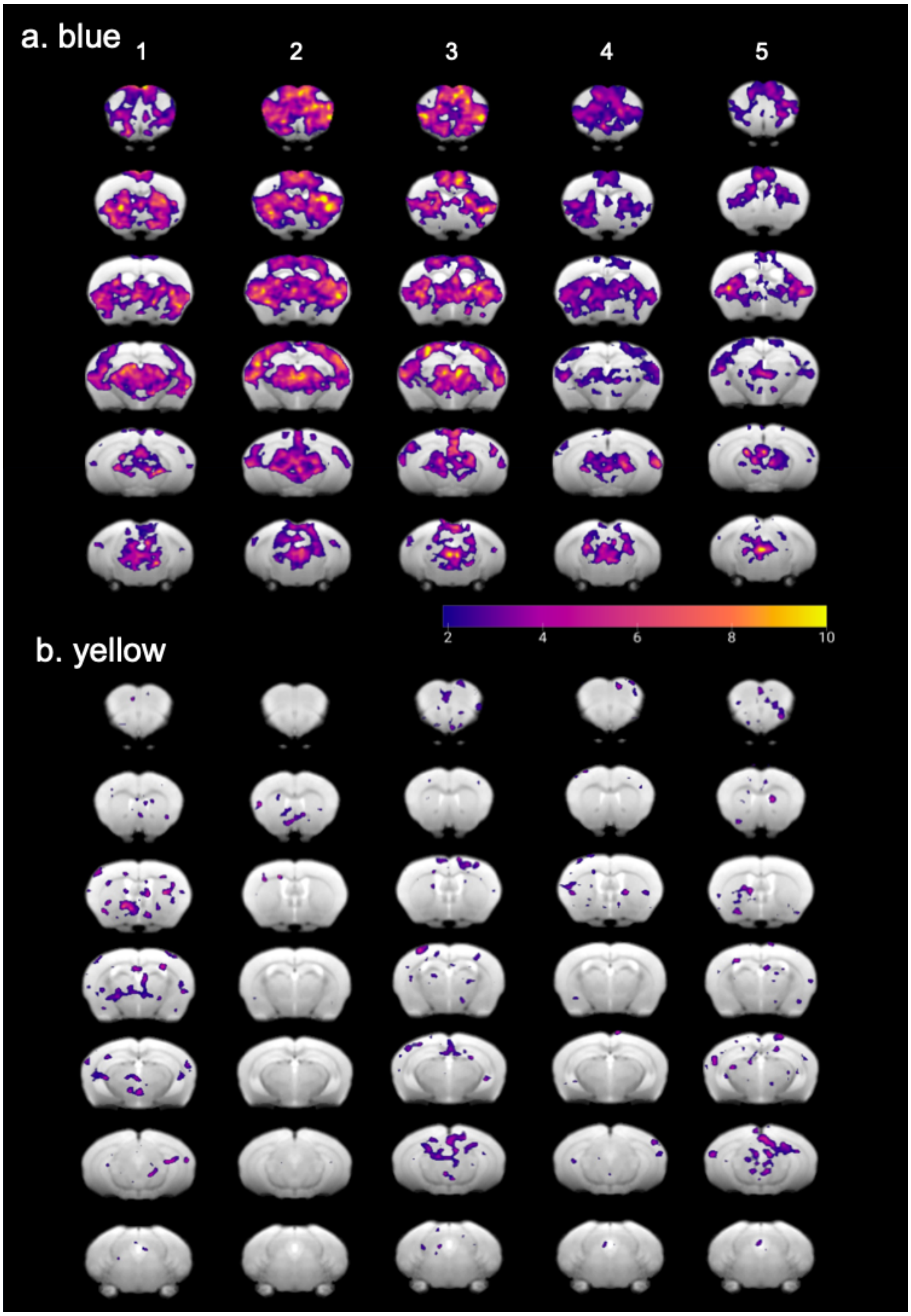


Supplementary Figure S2. Trial-wise brain response to optogenetic stimulations in Session 1. (a) Trial-wise functional maps of transgenic mice with blue illumination in Session 1 ($p_{uncorrected} < 0.05$, Transgenic: n = 8). The numbers 1-5 correspond to the timing of each stimulus. (b) Trial-wise functional maps of transgenic mice with yellow illumination in Session 1 ($p_{uncorrected} < 0.05$, Transgenic: n = 8).

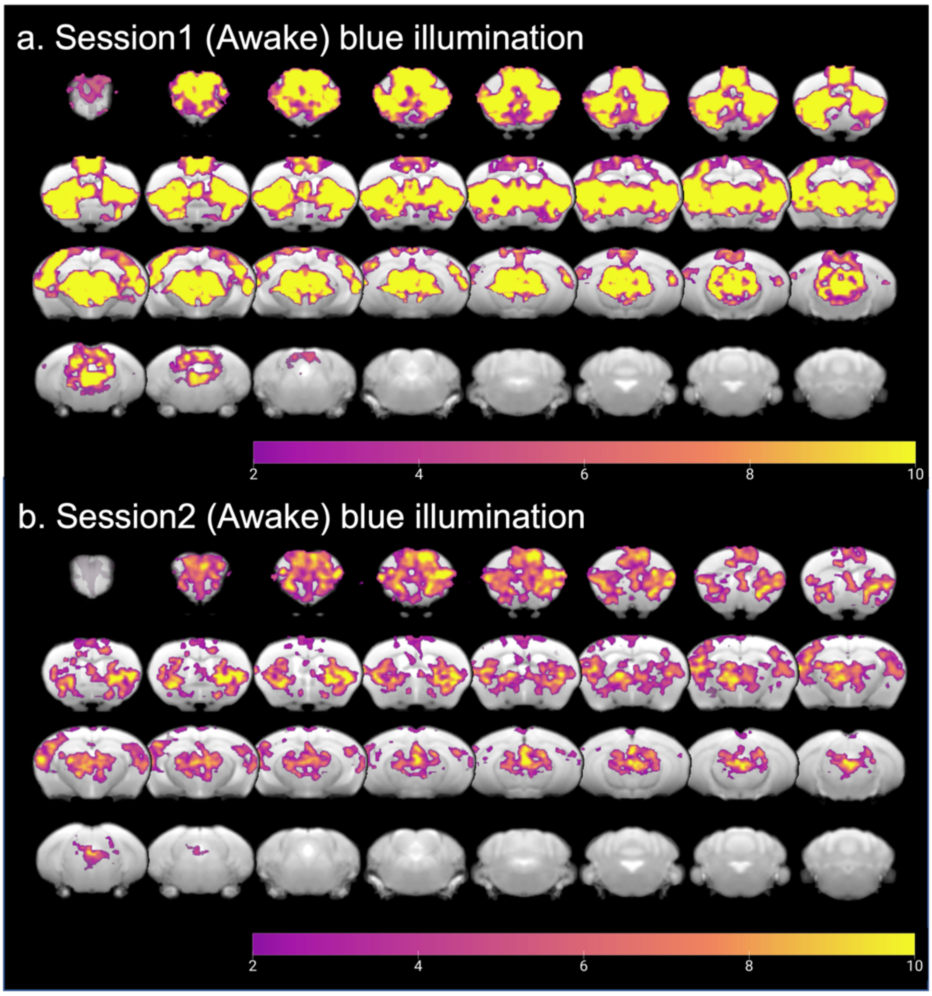

Supplementary Figure S3. Functional maps of transgenic mice in Session 1 and Session2 (a) Functional map of transgenic mice with blue illumination in Session 1 ($p_{uncorrected} < 0.05$, Transgenic: n = 8). (b) Functional map of transgenic mice with blue illumination in Session 2 ($p_{uncorrected} < 0.05$, Transgenic: n = 7).

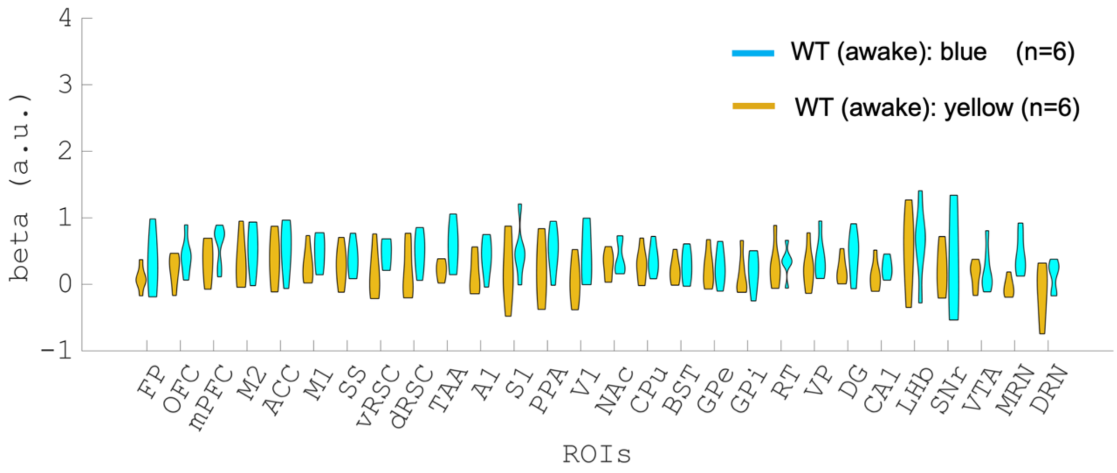


Supplementary Figure S4. Comparison of beta values in multiple ROIs in the WT group (n = 6). Beta values were obtained from GLM analysis in Session 1, and were extracted and averaged based on ROIs from the Allen Brain Atlas (Table1 and Supplementary Table ST2). The difference of beta values between blue and yellow illumination sessions were contrasted in the WT group. Cyan and orange colors indicate blue and yellow illumination trials, respectively.

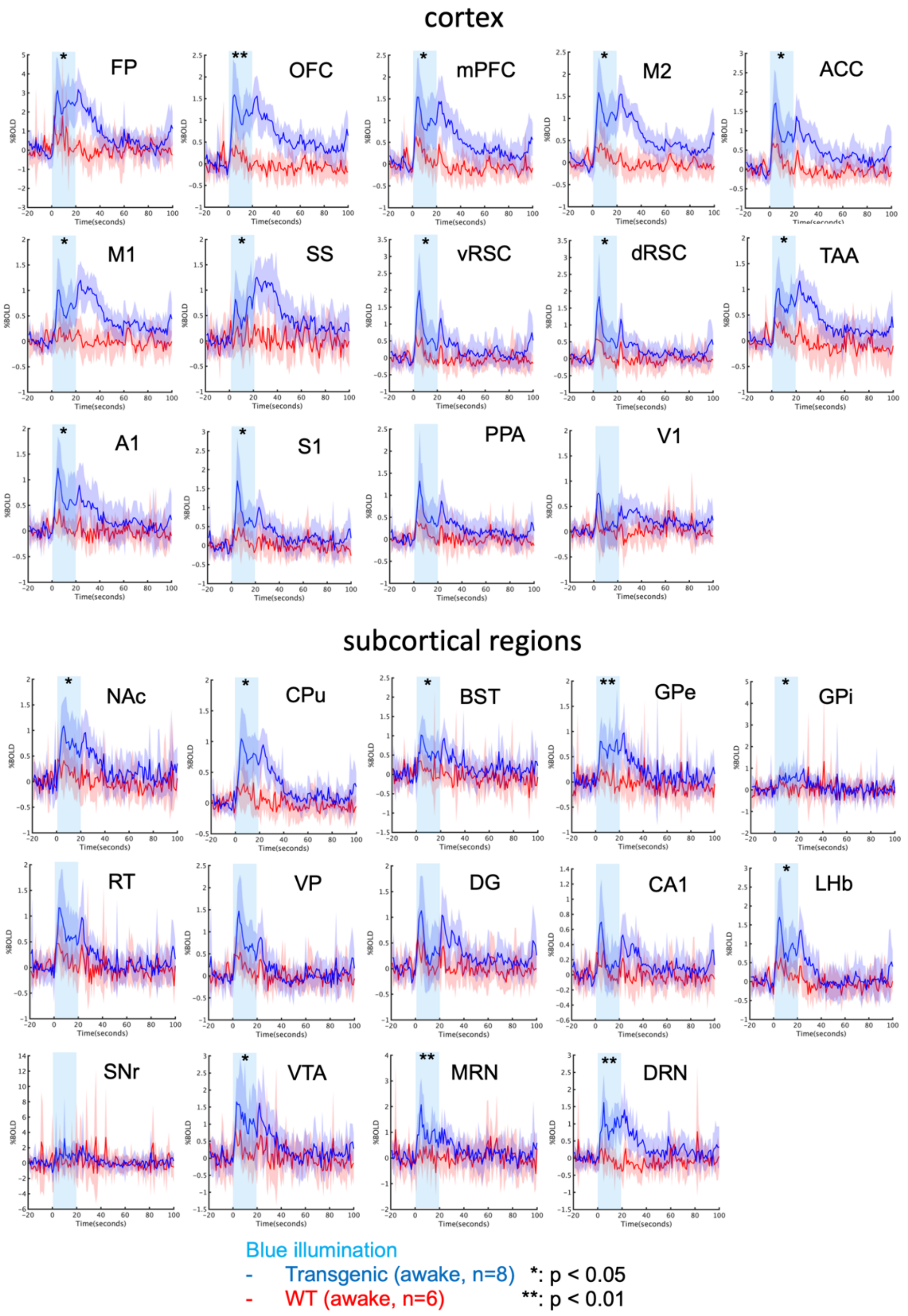


Supplementary Figure S5. Average % BOLD responses to blue illumination in Session 1. Time series of % BOLD responses to yellow illumination in each mouse were averaged at the onset of illumination. Average % BOLD responses were contrasted between the transgenic (n = 8) and WT groups (n = 6). Blue illumination (1.0 s; 473 nm) was applied at the onset of stimulation, and yellow illumination (1.0 second; 593 nm) followed the offset of stimulation 20 s after blue illumination. Cyan highlighting indicates the stimulation duration for 20 s. * and ** indicate $p_{FDR\text{-}corrected} < 0.05$ and 0.01, respectively (corrected with FDR).

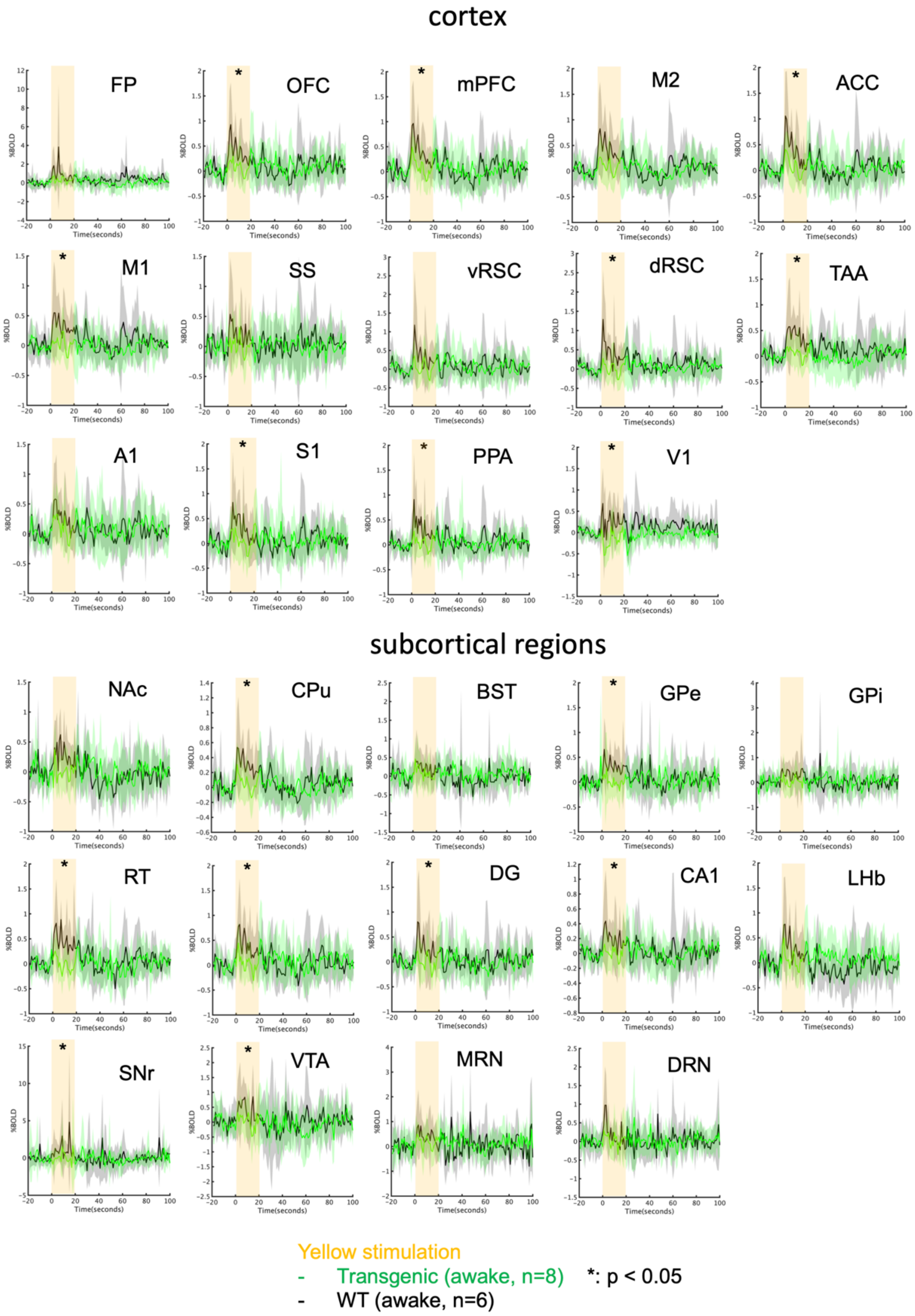


Supplementary Figure S6. Average % BOLD responses of yellow illumination in Session 1. Time series of % BOLD responses to yellow illumination in each mouse were averaged at the onset of illumination. Average % BOLD responses were contrasted between the transgenic (n = 8) and WT groups (n = 6). Yellow highlighting indicates the stimulus duration for 20 s. * indicate $p_{FDR} < 0.05$ (corrected with FDR).

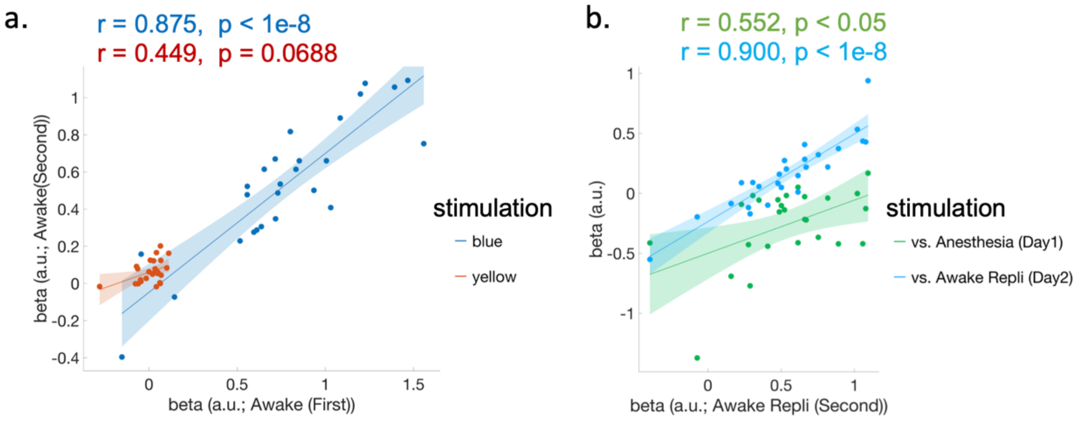


Supplementary Figure S7. BOLD response between Session 1-4. (a) Replication of average beta values from GLM to blue stimulation between Sessions 1 and 2 in transgenic mice. Statistically significant correlation was found in blue photo-stimulation ($r = 0.875$, $p_{uncorrected} < 1e\text{-}8$), but not in yellow photo-stimulation ($r = 0.449$, $p_{uncorrected} = 0.0688$). (b) Relationship of average regional beta values from GLM to blue photo-stimulation between Session 2 and Sessions 3 and 4 in transgenic mice. Statistically significant correlations were found in Session 3 under general anesthetic ($r = 0.552$, $p_{uncorrected} < 0.05$) and in Session 4 without general anesthesia ($r = 0.900$, $p_{uncorrected} < 1e^{-8}$).

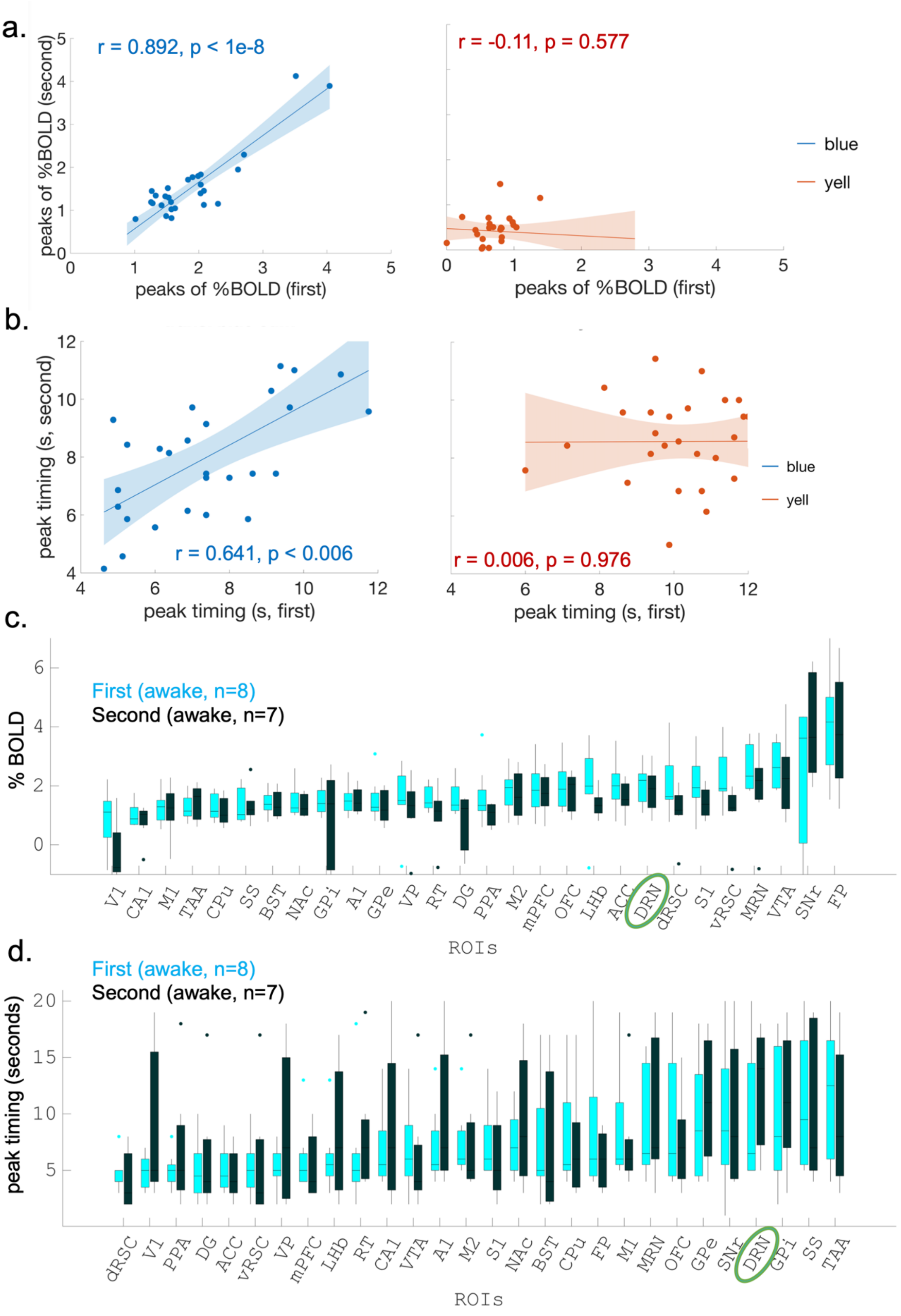


Supplementary Figure S8. Peak and timing of BOLD response in Session 1 and Session 2 (a) Replication of peaks of average % BOLD signals between Sessions 1 and 2 in transgenic mice. Statistically significant correlation between peaks of % BOLD responses to blue photo-stimulation in Sessions 1 and 2

were found, but not to yellow photo-stimulation (blue illumination: r = 0.8863, $p_{uncorrected} < 1e\text{-}8$; yellow photo stimulation: r = -0.06, $p_{uncorrected} = 0.766$). (b) Replication of peaks of average % BOLD signals between Sessions 1 and 2 in transgenic mice. Statistically significant correlation between peak timings of % BOLD responses to blue photo-stimulation in Sessions 1 and 2 was found, but not to yellow photo-stimulation (blue illumination: r = 0.6785, $p_{uncorrected} < 0.005$; yellow photo stimulation: r = 0.0369, $p_{uncorrected} = 0.8551$). (c) Alignment of peaks of regional % BOLD signals in Sessions 1 and 2. Peaks of % BOLD signals are sorted in ascending order. (d) Alignment of peak timing of regional % BOLD signals in Sessions 1 and 2. Peak timings of regional % BOLD signals are sorted in ascending order.

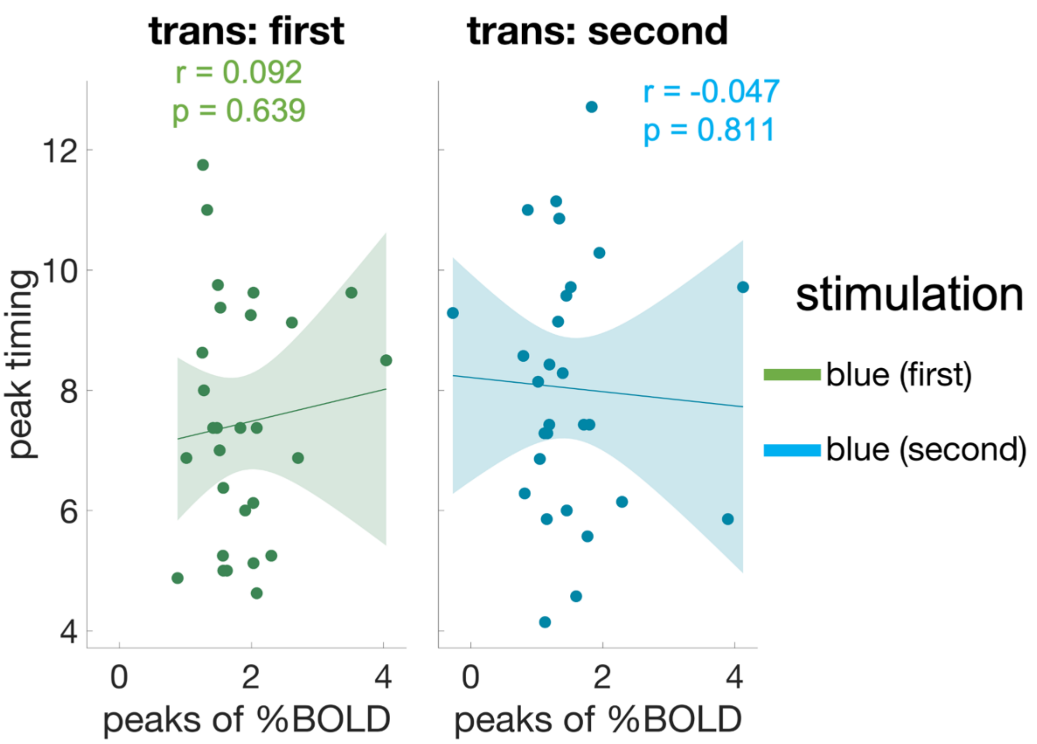


Supplementary Figure S9. Association between peak timing and peaks of average % BOLD signals in Sessions 1 and 2. No statistically significant correlations between peak timing and peaks of photo-stimulated %BOLD signals were found in either session (Session 1: $r = 0.092$, $p_{uncorrected} = 0.639$; Session 2: $r = -0.047$, $p_{uncorrected} = 0.811$).

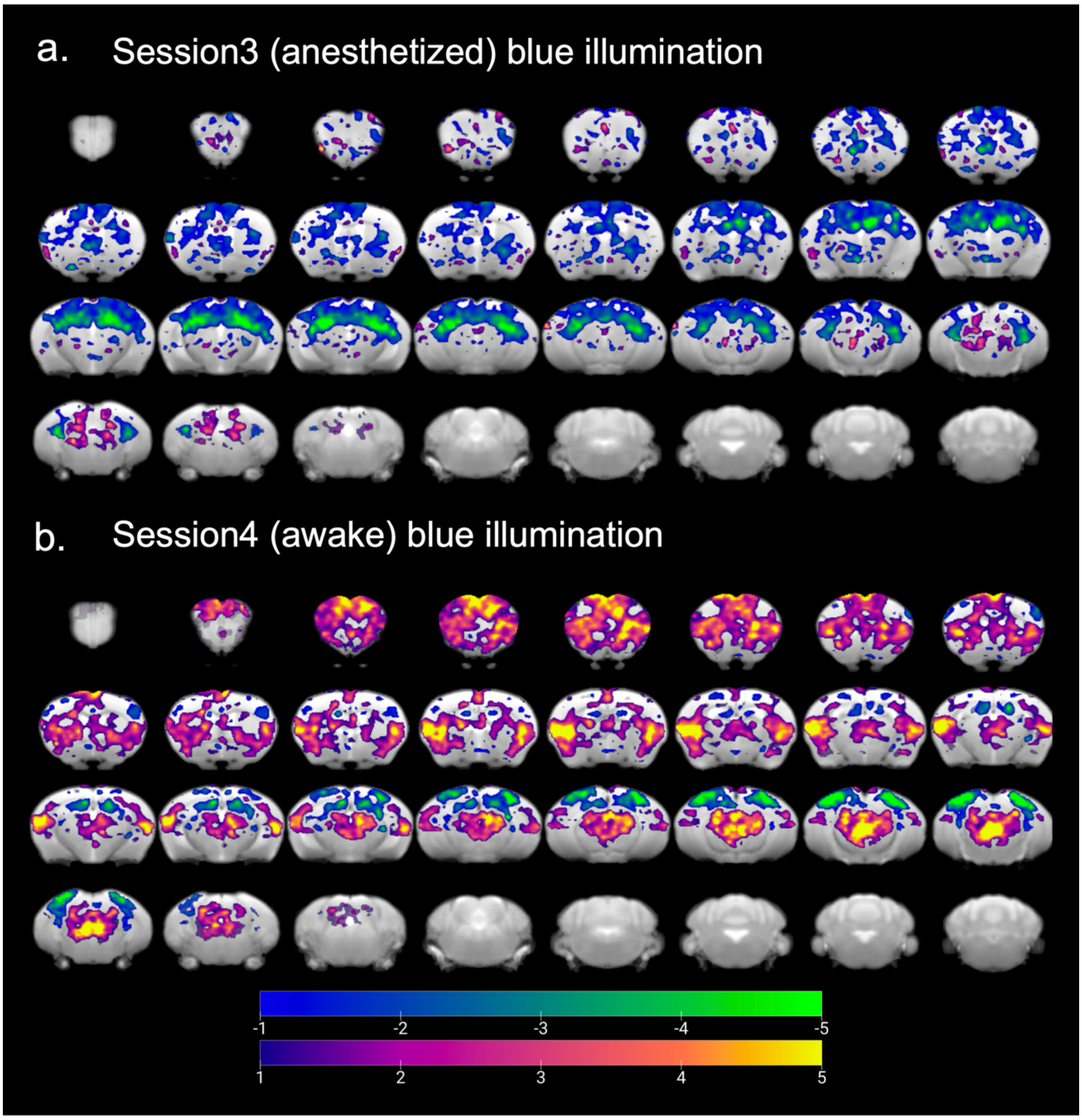


Supplementary Figure S10. Functional maps of transgenic mice in Session 3 and Session 4. (a) Functional maps of transgenic mice (n = 7) with blue illumination in Session 3 under anesthesia ($t_{uncorrected} > 1$). (b) Functional maps of transgenic mice (n = 7) with blue illumination in Session 4 without anesthesia ($t_{uncorrected} > 1$).

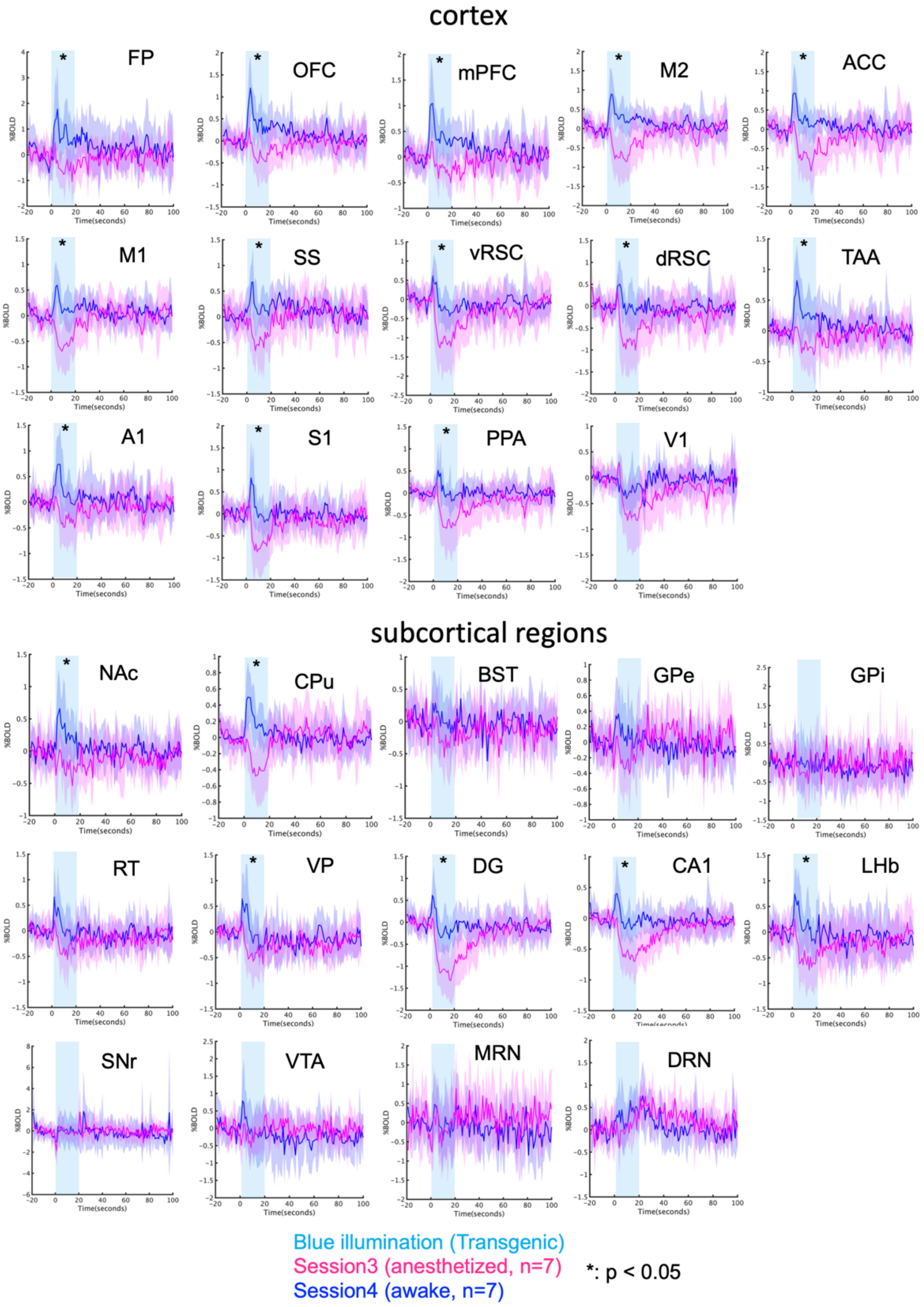


Supplementary Figure S11. Average % BOLD responses to blue illumination in Sessions 3 and 4. Time series of % BOLD responses by blue illumination in each mouse were averaged at the onset of illumination. Average % BOLD responses were contrasted between Sessions 3 and 4 in the transgenic group (n = 7). Blue illumination (1.0 s; 473 nm) was applied at the onset of stimulation. Yellow illumination (1.0 second; 593 nm) followed the offset of stimulation 20 s after blue illumination. Cyan highlighting indicates the stimulus duration for 20 s. * indicates $p_{FDR} < 0.05$ (corrected with FDR).

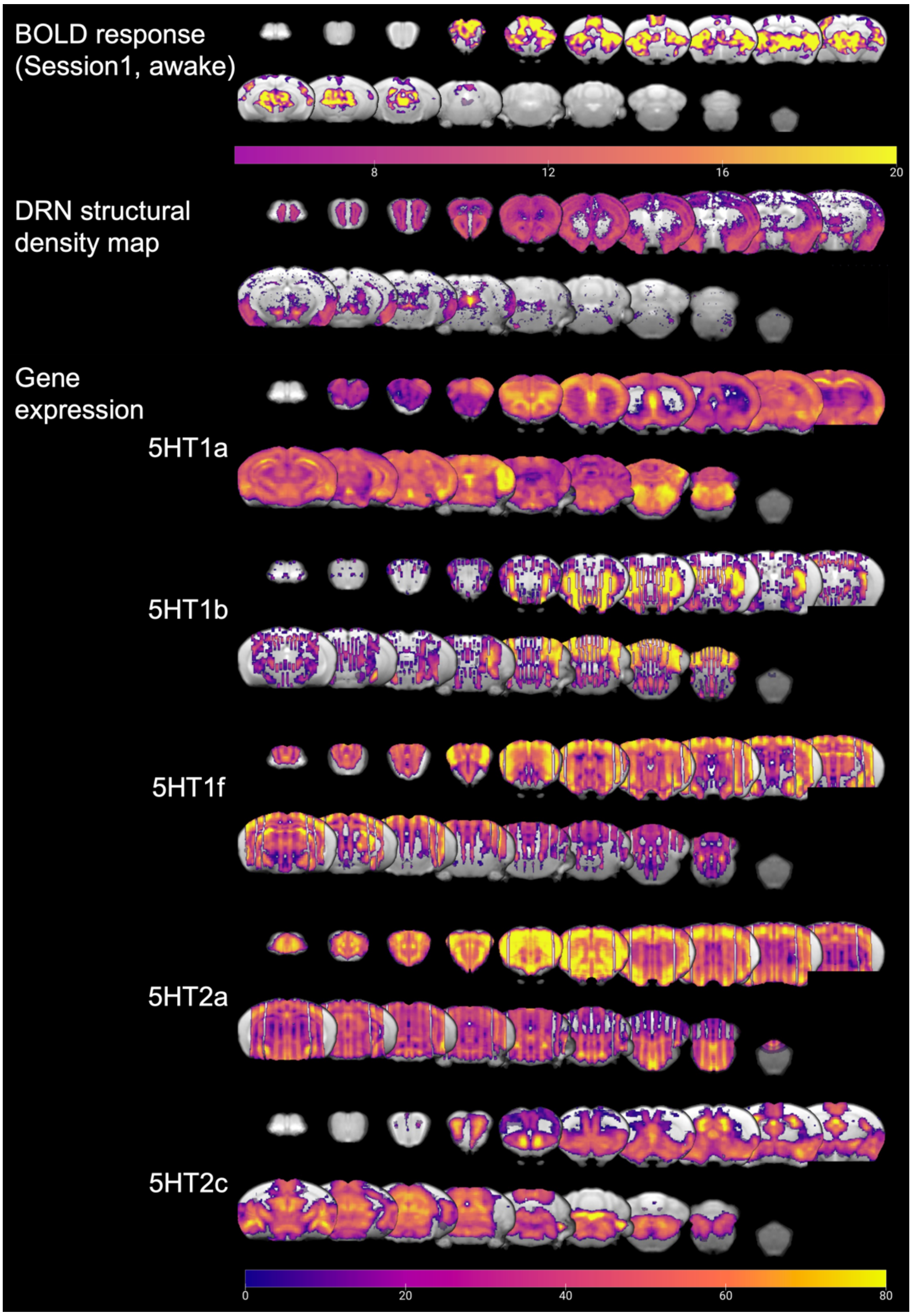


Supplementary Figure S12. Comparison of functional maps, structural density, and gene expression maps. The functional map from Session 1 under blue illumination was aligned with structural density and gene expression maps. The structural density map and gene expression of serotonin receptors are obtained from the mouse brain database of the Allen Brain Institute (Experimental IDs: 480074702 (5HT projection), 79556616 (5HT1a), 583 (5HT1b), 69859867 (5HT1f), 81671344 (5HT2a), and 73636098 (5HT2c)). A bottom color bar indicates normalized intensity of mRNA expression density of receptors and structural density for projections. The color bar ranges from 0-80.

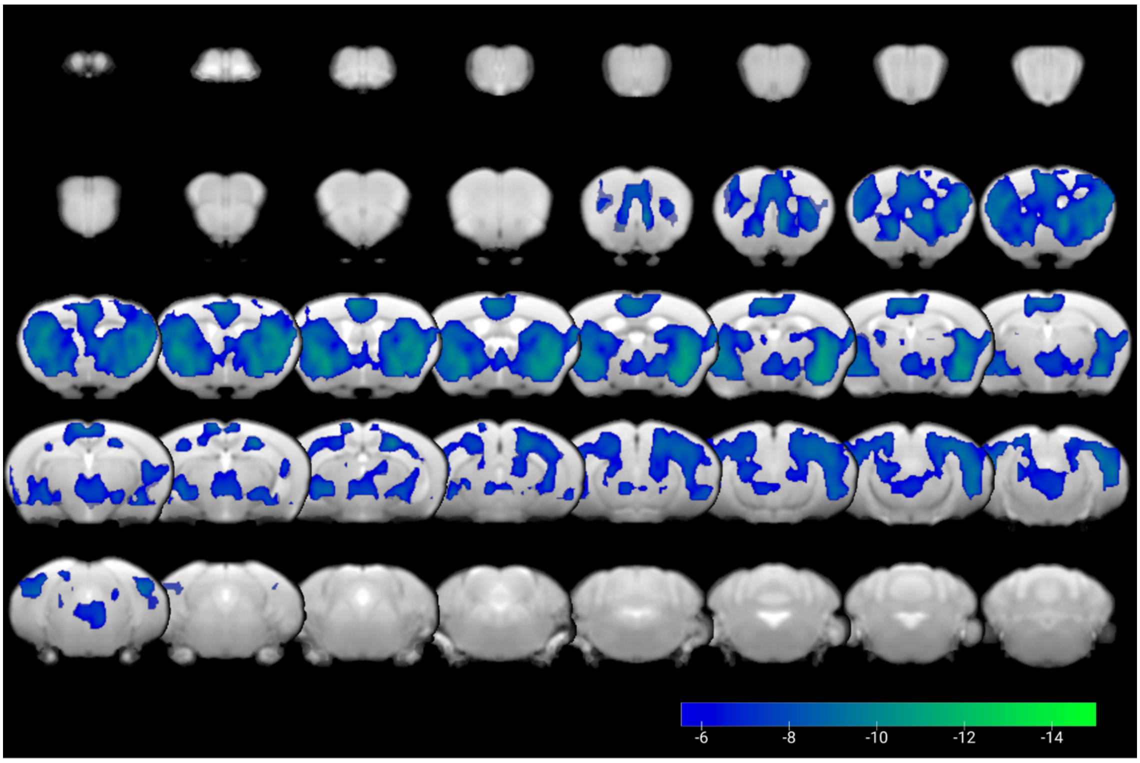


**Supplementary Figure S13.** Functional map of CBV response with blue illumination, from Grandjean et al., (2019) ($p_{FWE\text{-corrected}} < 0.05$, FWE-corrected; n=8, 63 runs). A bottom blue color bar indicates the statistical significance of the t value ($p_{FWE\text{-corrected}} < 0.05$). The color bar ranges from -5.6-15.

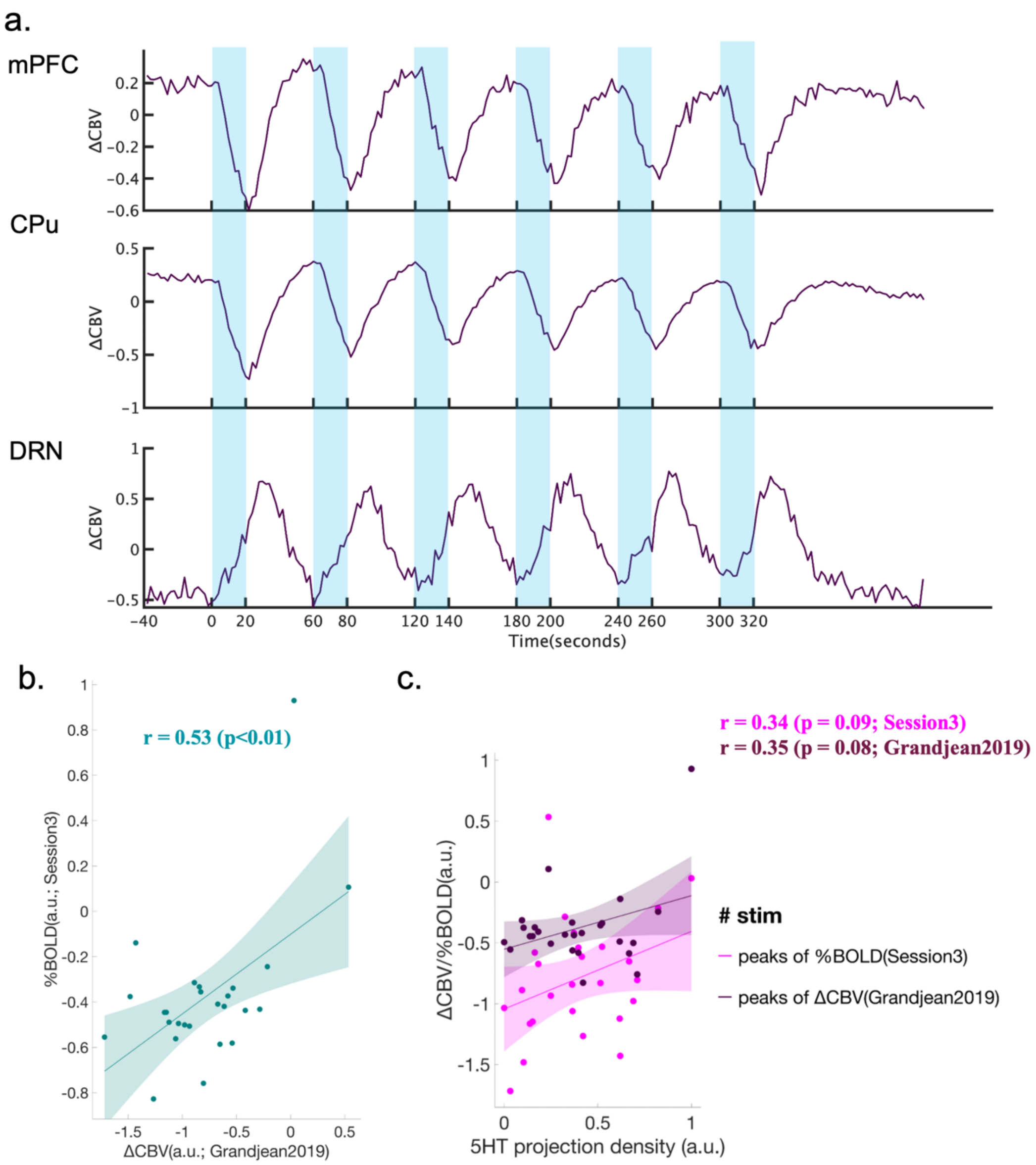


**Supplementary Figure S14.** Association of peak intensities between CBV signals [1, 3] and BOLD signals (Session 3). (a) Time series of regional CBV signals from mPFC, CPu, and DRN. Blue highlighting indicates 20-sec stimulation cycles. (b) Association of peak intensities between CBV signals and BOLD signals under anesthesia (Session3). Statistically significant correlation between peak intensities between the CBV signals and % BOLD signal was found ($r = 0.53$, $p_{uncorrected} < 0.01$). (c) Association between structural density and brain response. No statistically significant correlations of structural density with peak intensity of the CBV and % BOLD signals were found (vs. CBV signals (Grandjean2019): $r = 0.35$, $p_{uncorrected} = 0.08$; vs. %BOLD signals (Session3): $r = 0.34$, $p_{uncorrected} = 0.09$). Deep and light magenta indicate the association of structural density with the peak of %BOLD and that of ΔCBV, respectively.

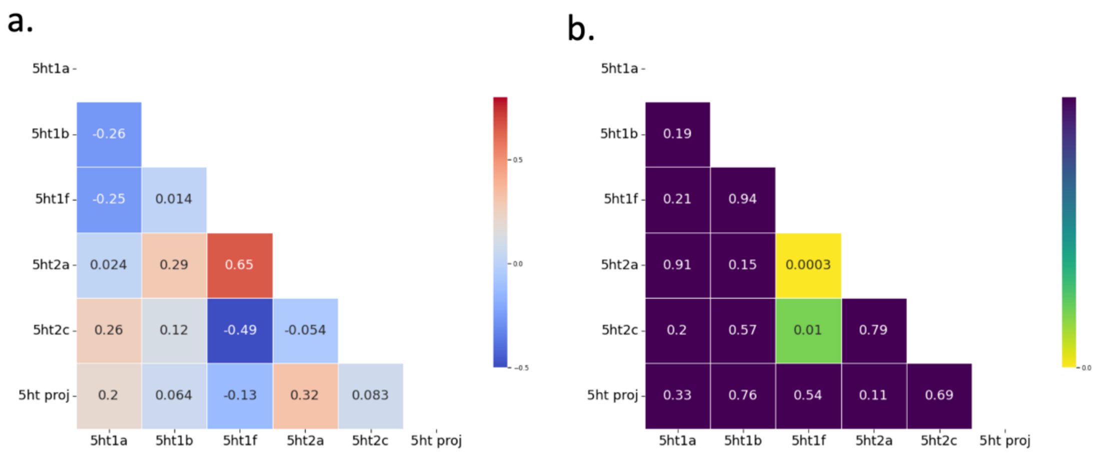


Supplementary Figure S15. Correlation between 5HT projection density and expression profiles of serotonin receptors. (a) Pearson's correlation of 5HT projection density and expression profiles of serotonin receptors. (b) Corresponding statistical p values of the correlation between 5HT projection density and expression profiles of serotonin receptors ($p_{uncorrected} < 0.05$, two-tailed t tests).

#### References


1. Grandjean, J., et al., *A brain-wide functional map of the serotonergic responses to acute stress and fluoxetine.* Nat Commun, 2019. **10**(1): p. 350.
2. Hikishima, K., et al., *In vivo microscopic voxel-based morphometry with a brain template to characterize strain-specific structures in the mouse brain.* Sci Rep, 2017. **7**(1): p. 85.
3. Mandino, F., et al., *A triple-network organization for the mouse brain.* Mol Psychiatry, 2022. **27**(2): p. 865-872.
4. Morel, P., *Gramm: grammar of graphics plotting in Matlab.* Journal of Open Source Software, 2018. **3**(23): p. 568.
5. Spisak, T., et al., *Probabilistic TFCE: A generalized combination of cluster size and voxel intensity to increase statistical power.* Neuroimage, 2019. **185**: p. 12-26.
